# Characterizations of enhanced infectivity and antibody evasion of Omicron BA.2.75

**DOI:** 10.1101/2022.07.18.500332

**Authors:** Yunlong Cao, Weiliang Song, Lei Wang, Pan Liu, Can Yue, Fanchong Jian, Yuanling Yu, Ayijiang Yisimayi, Peng Wang, Yao Wang, Qianhui Zhu, Jie Deng, Wangjun Fu, Lingling Yu, Na Zhang, Jing Wang, Tianhe Xiao, Ran An, Jing Wang, Lu Liu, Sijie Yang, Xiao Niu, Qingqing Gu, Fei Shao, Xiaohua Hao, Ronghua Jin, Youchun Wang, Xiaoliang Sunney Xie, Xiangxi Wang

**Author notes:** Correspondence: Yunlong Cao; Xiaoliang Sunney Xie; Xiangxi Wang. These authors contributed equally (Yunlong Cao, Weiliang Song, Lei Wang, Pan Liu, Can Yue, Fanchong Jian).

## Abstract

Recently emerged SARS-CoV-2 Omicron subvariant, BA.2.75, displayed a local growth advantage over BA.2.38, BA.2.76 and BA.5 in India. The underlying mechanism of BA.2.75’s enhanced infectivity, especially compared to BA.5, remains unclear. Here, we show that BA.2.75 exhibits substantially higher ACE2-binding affinity than BA.5. Also, BA.2.75 spike shows decreased thermostability and increased “up” RBD conformation in acidic conditions, suggesting enhanced low-pH-endosomal cell-entry pathway utilization. BA.2.75 is less humoral immune evasive than BA.4/BA.5 in BA.1/BA.2 breakthrough-infection convalescents; however, BA.2.75 shows heavier neutralization evasion in Delta breakthrough-infection convalescents. Importantly, plasma from BA.5 breakthrough infection exhibit significantly weaker neutralization against BA.2.75 than BA.5, mainly due to BA.2.75’s distinct RBD and NTD-targeting antibody escaping pattern from BA.4/BA.5. Additionally, Evusheld and Bebtelovimab remain effective against BA.2.75, and Sotrovimab recovered RBD-binding affinity. Together, our results suggest BA.2.75 may prevail after the global BA.4/BA.5 wave, and its increased receptor-binding capability could allow further incorporation of immune-evasive mutations.

## Introduction

Severe Acute Respiratory Coronavirus (SARS-CoV-2) Omicron variants have been continuously evolving and dominating the pandemic ^1,2^. Globally, BA.1 was rapidly replaced by the antigenically distinct descendant BA.2, while BA.4, BA.5 and BA.2.12.1, derived from BA.2, exhibited further increased humoral immunity evasion and have outcompeted BA.2 ^3–5^. More recently, a novel BA.2 subvariant designated as BA.2.75, a variant of concern (VOC) lineage under monitoring by the World Health Organization (WHO), is spreading rapidly in India and around the globe, contributing to over 20% of recently reported sequences in India and is continuously increasing ^1,6^. Compared to the BA.2 spike (S-trimer), BA.2.75 carries nine additional mutations, among which five are on the N-terminal domain (NTD), including K147E, W152R, F157L, I210V and G257S, and four on the receptor binding domain (RBD), namely D339H, G446S, N460K and R493Q (Figure 1A and 1B). Among them, G446S appeared in BA.1, and R493Q reversion was observed in BA.4/BA.5. N460K and D339H mutations have not been observed on prevailing variants, and their functions remain unclear.

**Fig. 1.**
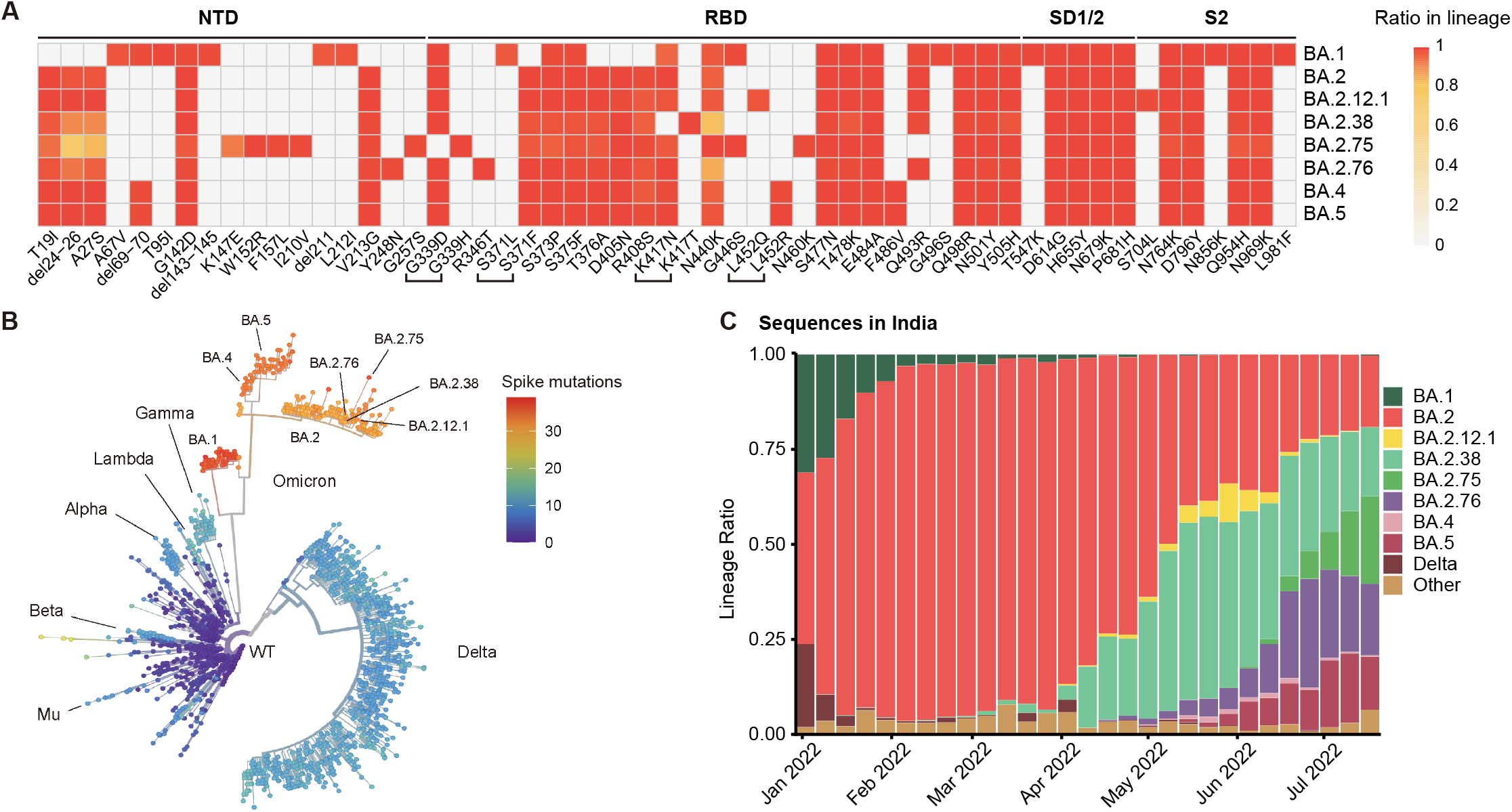
BA.2.75 and BA.2.76 displayed growth advantage in India. (A) Main mutations on the spike glycoprotein appearing in SARS-CoV-2 Omicron sublineages. (B) Phylogenetic tree of existing SARS-CoV-2 variants. Color scales indicate number of mutations on the spike. (C) Lineage distribution of recent sequences from India by time. BA.2.75 and BA.2.76 is growing rapidly in India, showing advantage compared to other lineages.

Importantly, BA.2.75 displayed a local growth advantage in India compared to BA.2.38 (BA.2+N417T), BA.2.76 (BA.2+R346T+Y248N) and BA.5 (Figure 1C). The enhanced transmissibility over BA.5 questions whether BA.2.75 would prevail after the global BA.4/BA.5 wave. To address this, the receptor-binding affinity and humoral immune evasion capability of BA.2.75, especially under the immune background after BA.4/BA.5 infection, need immediate evaluation.

## Results

### BA.2.75 displays higher ACE-binding affinity than BA.4/BA.5

Recent studies have revealed that Omicron subvariants have further improved binding affinities for hACE2 ^3–5,7^ and stability *in vitro*, which, to some extent, correlates with their increased viral transmission and infection. The difference in amino acid composition of the RBD between BA.2 and BA.2.75 is four substitutions: D339H, G446S, N460K and R493Q, among which G446S and R493Q lie in the RBD/hACE2 interface, N460K and D339H locate close to and far away from the interface, respectively (Figure 2A). To explore the impact of these mutations on the human ACE2 (hACE2) binding of BA.2.75 RBD, we assessed the binding affinities between hACE2 and RBDs from the six Omicron subvariants (BA.1, BA.2, BA.3, BA.4/5, BA.2.12.1 and BA.2.75), together with the other earlier four VOCs (Alpha, Beta, Gamma, and Delta) by surface plasmon resonance (SPR) (Figure 2B and Figure S1A). Surprisingly, BA.2.75 displayed a 3-6-fold increased binding affinity to hACE2 compared to other Omicron variants, substantially higher than the other four VOCs as well, reaching the highest binding activity measured in circulating SARS-CoV-2 strains so far.

**Fig. 2.**
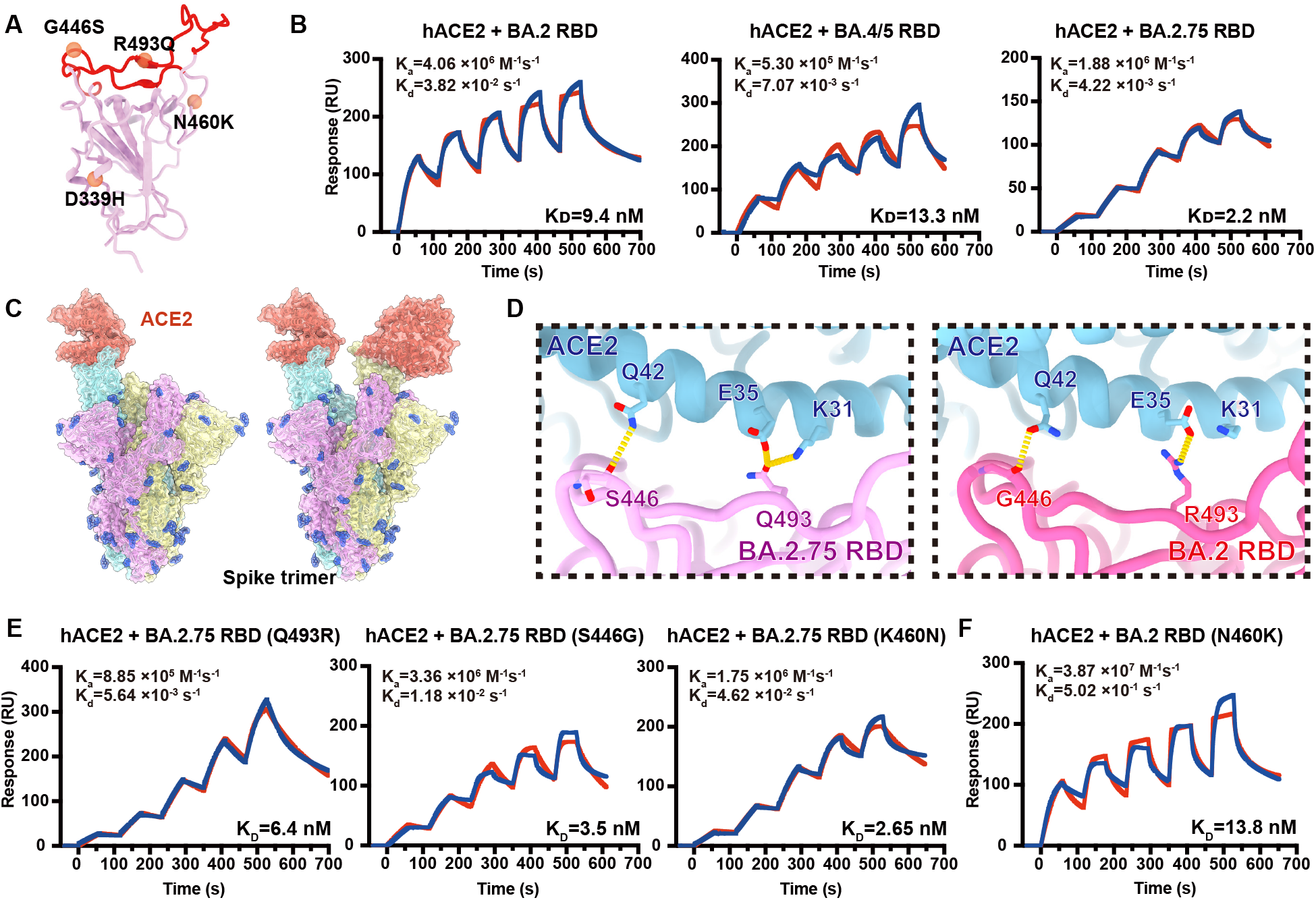
BA.2.75 exhibited enhanced human ACE2 binding. (A) Position distribution of important amino acids in BA.2.75 RBD. The mutated residues relative to BA.2 RBD are marked as orange globules. RBM is colored in red. (B) Binding affinities of RBDs of BA.2, BA.4/5 and BA.2.75 subvariants to hACE2 measured by SPR. (C) Overall structure of the BA.2.75 S-trimer in complex with hACE2. Three copies of S monomer were colored in yellow, cyan, and magenta, respectively. The hACE2 molecules bound to RBD were colored in orange. (D) Changes at the interfaces between BA.2.75 RBD (left) and BA.2 RBD (PDB: 7ZF7, right) with hACE2. Key mutated residues are shown as sticks and hydrogen bonds are shown as yellow dash lines. (E) Binding affinity of hACE2 with BA.2.75 RBD with single substitution Q493R, S446G, K460N measured by SPR. (F) Binding affinity of hACE2 with BA.2+N460K RBD measured by SPR.

To further unveil the molecular details of the enhanced hACE2 binding, we determined the near-atomic structure of the BA.2.75 S-trimer in complex with hACE2 (Figure 2C, Figure S1B and Table S1). Like most complex structures, two copies of hACE2 are bound to the two RBDs in the up conformation (Figure 2C). Structural comparisons revealed that the substitution of Q493R in BA.2.75 established one extra hydrogen bond with K31 on ACE2, increasing its binding capability (Figure 2D) ^4,8^. By contrast, the hydrophilic interaction between the main-chain carbonyl group of either G446 or S446 with Q42 from hACE2 is retained (Figure 2D), and mutations of N460K and D339H cause no notable alterations in the binding interface.

To further deconvolute the contribution on hACE2 binding of each mutation on BA.2.75 RBD, we evaluated the individual reverse mutation of H339D, S446G, K460N or Q493R in BA.2.75 RBD on the ACE2 affinity. SPR assays revealed that only Q493R site mutation induces an approximately 3-fold affinity decrease, while the other three reversions did not substantially alter the binding affinity (Figure 2E and S1C). Notably, previous deep mutational scanning results suggested that N460K would slightly enhance the ACE2-binding affinity in BA.2 background ^9^. To further confirm whether N460K could impact ACE2 binding, we measured the hACE2-binding affinity of BA.2+N460K RBD. Results demonstrate that N460K does not significantly affect hACE2-binding in both BA.2 and BA.2.75 background (Fig. 2F).

### Structural analyses of BA.2.75 spike suggest enhanced endosomal pathway usage

Omicron can enter into host cells endosomally as well as through TMPRSS2, but prefers endosomal fusion. TMPRSS2-dependent cell-surface fusion involves engaging at neutral pH, while the endosomal pathway proceeds with acidic pH ^10–12^. To investigate the putative alterations in cell-entry properties of BA.2.75, We determined the asymmetric cryo-EM reconstructions of the BA.2.75 S-trimer at an overall resolution of 2.8-3.5 Å at serological (pH 7.4) and endosomal pHs (pH 5.5) (Fig. 3A, Fig. S2A-B, and Table S1). Like BA.2 and BA.2.12.1, the BA.2.75 S-trimer exhibit two distinct conformational states (mol ratio ≈ 1:1) corresponding to a closed-form with all three RBDs “down” and an open form with one RBD “up” at neutral pH (Fig. 3A). Interestingly, single-RBD-up conformations dramatically dominated at pH 5.5 with mol ratio of 3:1. In addition, multiple orientations of RBD in the S-trimer were observed at pH 5.5, revealing structural heterogeneity in BA.2.75, akin to structural observations in D614G and Delta ^12^, suggesting putative enhanced viral fusion efficiency (Fig. 3B-C). *In vitro* thermal-stability evaluation indicated that the BA.2.75 S-trimer was the most stable among Omicron variants with a melting temperature of 66 °C at neutral pH, 3 °C improved compared to BA.1 (Fig. 3D). Unlike the other four subvariants, BA.2.75 exhibited a distinct shift in stability, less stable than BA.1 at endosomal pH. Together, results suggest BA.2.75 may have evolved to further utilize the low-pH-endosomal cell-entry pathway.

**Fig. 3.**
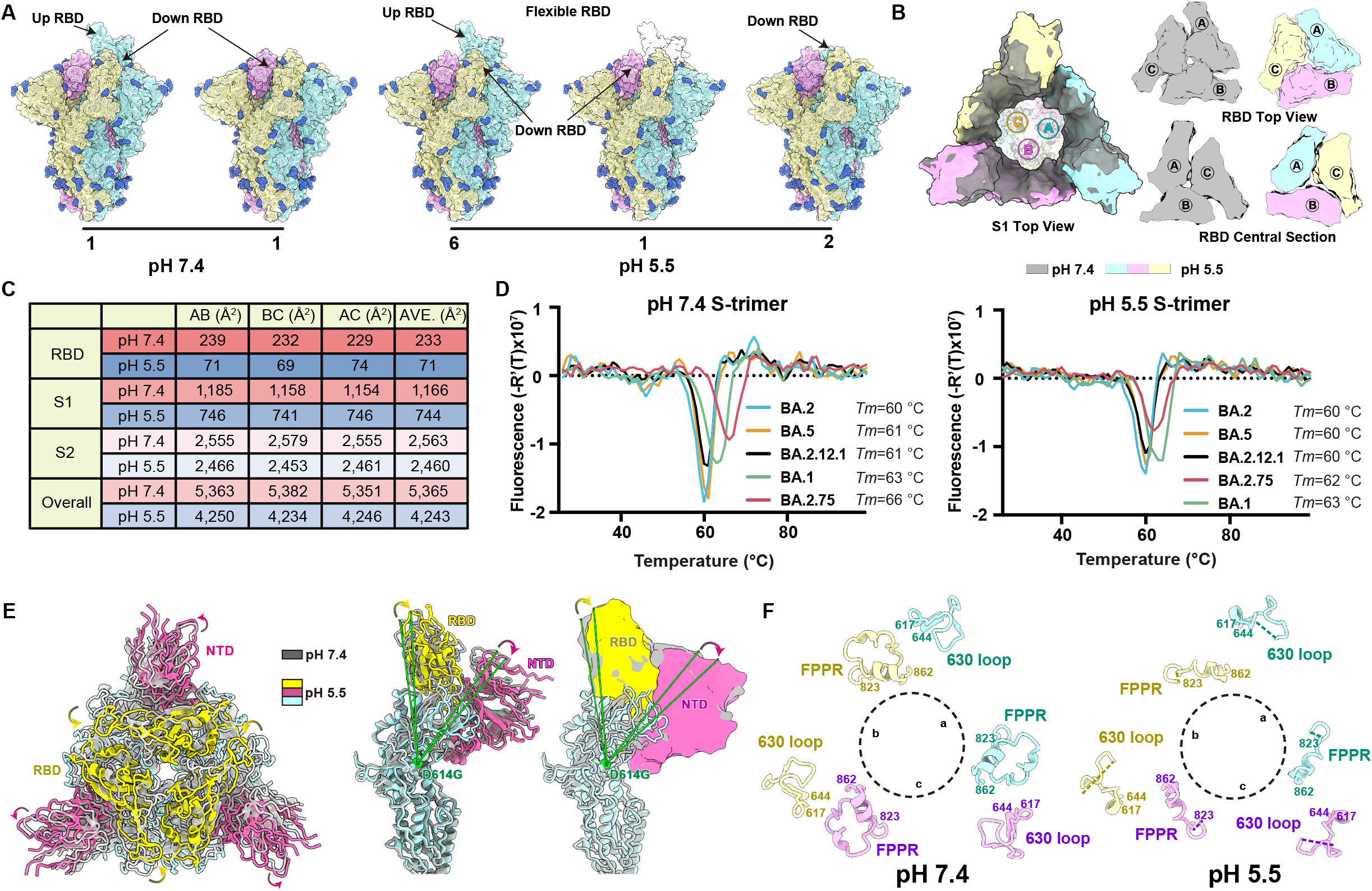
Structural characteristics of BA.2.75 spike glycoprotein. (A) Surface representation for structures of S-trimer of BA.2.75 at neutral (pH=7.4) and acidic conditions (pH=5.5); three protomers were colored in yellow, light blue and pink, respectively and N-glycans were highlighted in deep blue. (B) Structural organization of three S1-subunits and RBDs from the neutral (gray) and acidic BA.2.75 S-trimer (yellow, light blue and pink). Top view of S2 subunit (left), top view of the RBD (upper right) and central section of RBD (bottom right) show the inter-subunit contacts of the BA.2.75 S-trimers in different pH. (C) Buried surface areas between two neighboring protomers, S2-subunits, S1-subunits or RBD domains. (D) Thermal stability analysis of BA.1 (green), BA.2 (blue), BA.2.12.1 (black), BA.2.75 (red) and BA.5 (yellow) S-trimers at neutral and acidic pHs. (E) Superimposition of the neutral BA.2.75 S-trimer structure (grey) onto the structure of the acidic BA.2.75 S-trimer (RBD, yellow; NTD, hot pink); Structural rotations and shifts between these two structures were marked by green lines and arrows. (F) Structural alterations in the 630 loop (residues 617 to 644, light blue) and FPPR (residues 823 to 862, yellow) of the three protomers (a, b, and c) from these two S-trimers were shown. Dashed lines indicate gaps in the chain trace (disordered loops).

Compared to other Omicron subvariants, increased structural heterogeneity in BA.2.75 is mainly contributed by the reduced hydrophilic and hydrophobic interactions between two contacted RBDs due to the reversion of R493Q and local conformational alterations in the hairpin loop (residues 373-380), respectively (Fig. S2C). Surprisingly, the BA.2.75 RBD displays one more rigid and compact configuration than other subvariants, presumably representing improved stability and immunogenicity (Fig. 4A). Substitution of N460K established one new salt bridge with D420 and mutation of D339H with altered rotamer formed π-π interactions with F371, pulling α1 and α2 helices closer compared to BA.2 and BA.1, respectively (Fig. 4A and Fig. S3). Perhaps correlated with this, molecular simulation analysis also revealed overall improved stability for the RBD and the distal RBM in BA.2.75, which might facilitate the protein folding and increase protein expression (Fig. 4B). These structural analyses are in line with experimental observations revealed by thermal stability and deep mutational scanning results of N460K/D339H (Fig. 4C) ^9,13,14^.

**Fig. 4.**
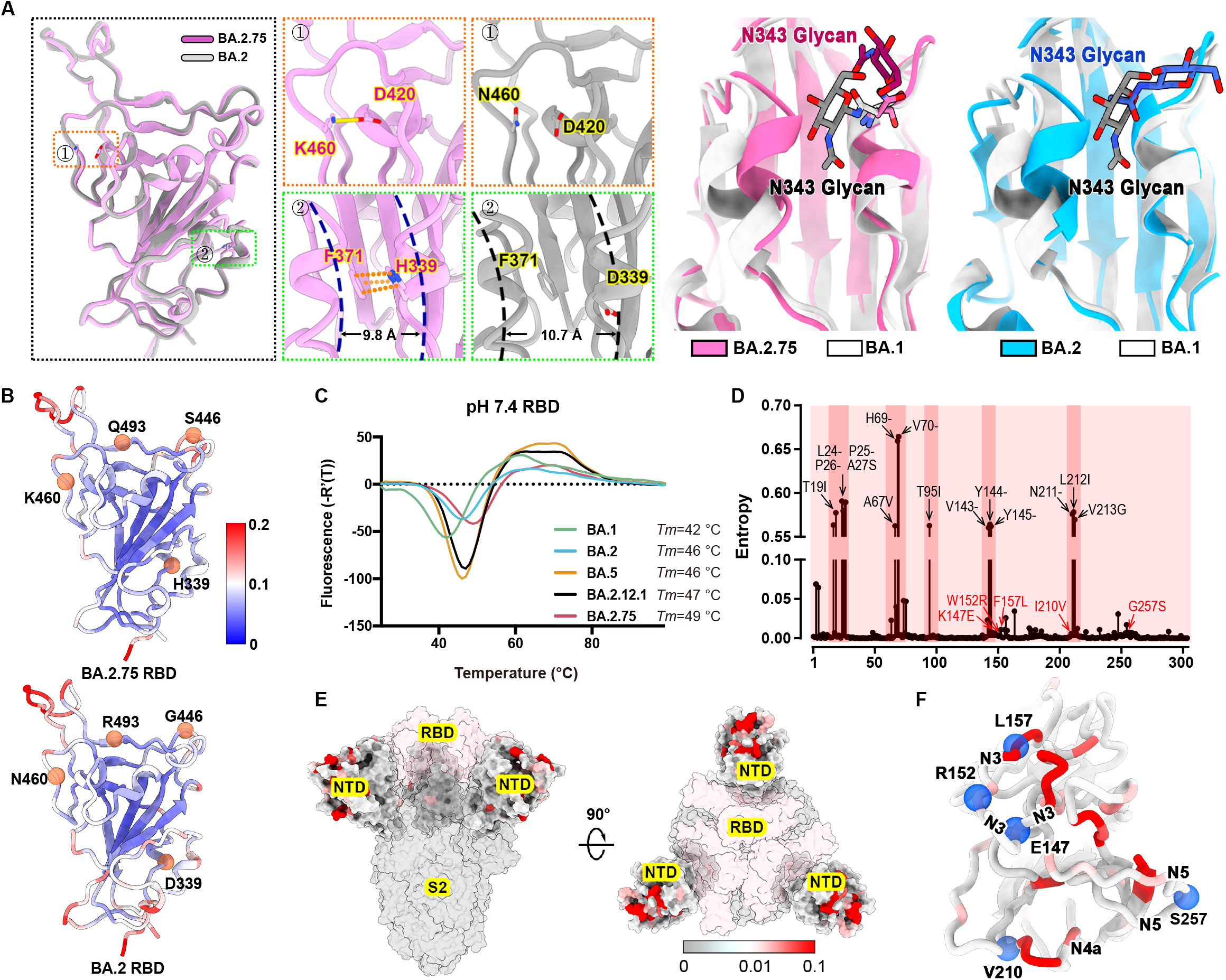
Structural features of BA.2.75 spike RBD and NTD. (A) Structural comparisons of RBDs of BA.2.75, BA.2 and BA.1. The newly established interaction of BA.2.75 (pink) with respect to BA.2 (gray) on the RBD is shown on the left. Salt Bridges formed between D420 and K460 and π-π stack formed between H339 and F371 in BA.2.75 RBD are highlighted. The distances between α1 and α2 helices on RBD are also marked. A diagram presentation of N343 glycan conformational differences among BA.1 (gray), BA.2 (blue), and BA.2.75 (pink) is shown on the right. (B) The stability landscapes of BA.2.75 and BA.2 RBD calculated by MD stimulation. The cartoons of BA.2.75 and BA.2 RBD are colored by root mean square fluctuation (RMSF) calculated from the last 2 ns of the stimulations. Residues 339, 446, 460 and 493 are shown as red spheres. (C). Thermal stability measurements of the RBD from BA.1 (green), BA.2 (blue), BA.5 (orange), BA.2.12.1 (black) and BA.2.75 (red) at pH 7.4. (D). Entropy of SARS-CoV-2 NTD variants among circulating isolates. The residues with higher entropy are highlighted by dark red background. The dominant mutations on SARS-CoV-2 NTD and the mutations on BA.2.75 NTD are labeled in black and red, respectively. (E). The heatmap for circulating variants with mutations on the NTD. Mutation frequency for each residue was calculated based on the datasets from GISAID.

The NTD shows the most diversity with a larger number of prevalent mutations and deletions compared to other regions of the S-trimer (Fig. 4D). Interestingly, the majority of these substitutions are highly enriched in the peripheral region of the S-trimer, adjacent to the NTD super-site (Fig. 4E), presumably escaping neutralizing antibodies and modulating entry efficiency ^15,16^. Among five mutations in the BA.2.75 NTD, K147E, W152R, and F157L locate at the super-site, while the other two partially overlap with epitopes targeted by other neutralizing antibody classes (Fig. 4F).

### BA.2.75 significantly evades plasma from Delta and BA.4/BA.5 convalescents

Next, we evaluated the effect of BA.2.75 on the neutralizing activities of vaccinees/convalescents plasma. Pseudovirus neutralization assays show that BA.2.75 exhibits significantly stronger humoral immune evasion than BA.2 in plasma samples from individuals who had received 3 doses of CoronaVac before or after BA.1/BA.2 breakthrough infection, resulting in a 1.4 to 3.2-fold reduction in half neutralization titers (NT50) (Fig. 5A and Table S2). BA.2.75 is also slightly more neutralization evasive than BA.2.12.1 in post-vaccination BA.2 convalescents, comparable to BA.2.76 but less than BA.4/BA.5 (Fig. 5A). However, BA.2.75 is more humoral immune evasive than BA.4/BA.5 in plasma from Delta breakthrough-infection convalescents, which may explain BA.2.75’s substantial growth advantage in India (Fig. 5A). This phenomenon could be contributed by the R452 stimulation of Delta, which could result in potent antibodies that are highly effective against BA.4/BA.5. Most importantly, BA.2.75 also displays strong humoral immune evasion in the convalescent plasma from BA.5 breakthrough-infection convalescents (Fig. 5A). Compared to BA.5, the NT50 against BA.2.75 by BA.5 convalescent plasma showed a nearly 4-fold reduction. Considering these samples also showed significantly decreased neutralization against BA.1, G446S should contribute majorly.

**Fig. 5.**
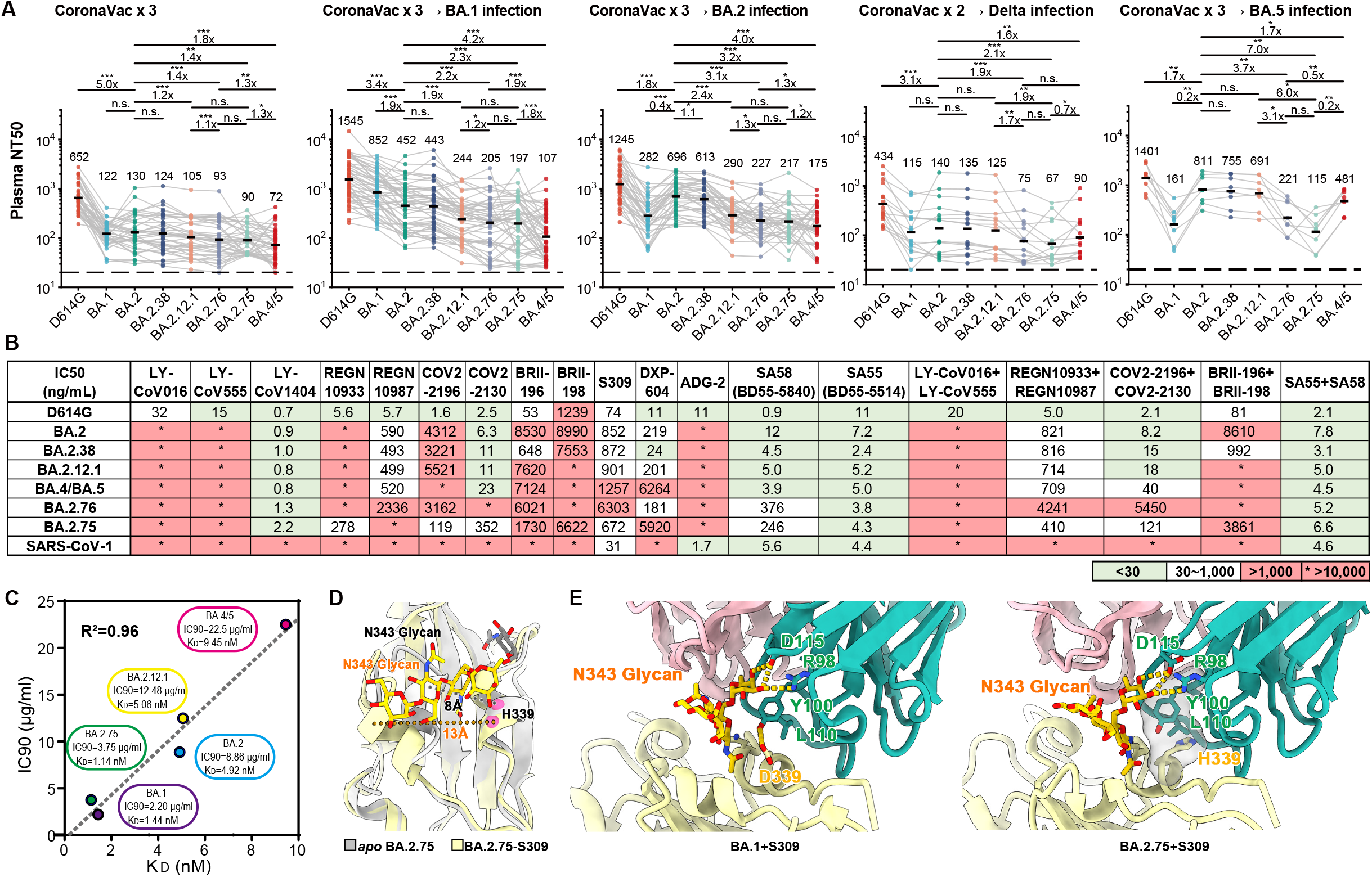
BA.2.38, BA.2.75 and BA.2.76 showed distinct antibody evasion. (A)Half neutralization titers (NT50) against SARS-CoV-2 D614G and Omicron variants pseudoviruses by plasma samples from individuals who received 3 doses CoronaVac (N=40), 3 doses CoronaVac followed by BA.1 infection (N=50), 3 doses CoronaVac followed by BA.2 infection (N=39), 2 doses CoronaVac followed by Delta infection (N=16), or 3 doses CoronaVac followed by BA.5 (N=8) infection. Geometric mean titers (GMT) were annotated above each group. *p < 0.05; **p < 0.01; ***p<0.001. P-values were calculated using two-tailed Wilcoxon singed-rand test of paired samples. (B) Neutralizing activities against SARS-CoV-2 D614G, Omicron variants, and SARS-CoV-1 pseudovirus of therapeutic neutralizing antibodies. Background colors indicate neutralization levels. Green, IC50 < 30 ng/mL; white, 30 ng/mL < IC50 < 1,000 ng/mL; red, IC50 > 1,000 ng/mL. *, IC50 > 10, 000 ng/mL. (C)Correlation plots of binding affinities and neutralizing activities (IC90) of S309 against BA.1, BA,2, BA.2.12.1, BA.4/5 and BA.2.75. (D)Local conformational alterations in the BA.2.75 RBD upon S309 binding. The N343 glycan of the apo BA.2.75 RBD (gray) and S309-bounded BA.2.75 RBD (yellow) are shown as sticks and the distances between α1 and α2 helices from two configurations are also labeled. (E)Interaction details of the BA.1 RBD (left) and BA.2.75 RBD (right) in complex with S309. Hydrogen bonds and hydro-phobic patches are presented as yellow dashed lines and gray surfaces, respectively. The light chain and heavy chain of S309 are colored in pink and cyan, respectively.

### Efficacy of NAb drugs against BA.2.75

As for antibody therapeutics, we tested the pseudovirus neutralizing activity of 14 NAb drugs in clinical development against SARS-CoV-2 variants, including BA.2.38, BA.2.75 and BA.2.76 ^17–25^. REGN10933 and COV2-2196 partially recovered their activities against BA.2.75 due to R493Q reversion (Fig. 5B). However, REGN10987 and COV2-2130 were also affected by G446S, resulting in only a mild change in the neutralizing activity of the corresponding cocktails against BA.2.75 (Fig. 5B) ^19,23,26^. LY-CoV1404 (bebtelovimab) remains highly potent against BA.2.75 and BA.2.76 ^17^.

Notably, multiple studies have reported the efficacy of S309 (Sotrovimab^18^) against BA.2.75; however, some suggest S309 recovered potency against BA.2.75, while others suggest the opposite ^27–29^. We noticed that although D339H is a charge-reversing mutation on the S309 binding interface, the neutralizing activity of S309 was not affected and even exhibited slightly improved neutralization activity against BA.2.75 compared to BA.2 and BA.4/5 (Fig. 5B). To further confirm this observation, we measured the binding affinity of S309 against BA.1, BA.2, BA.2.12.1, BA.4/5, and BA.2.75 RBD using biolayer interferometry (BLI). Interestingly, we found that S309 recovered its binding activity against BA.2.75 RBD as to that against BA.1, and the binding affinities of S309 to these five Omicron subvariants matched well with the IC90s, but not the IC50s (Fig. 5C and Fig. S4). To explore the underlying mechanism, we determined the cryo-EM structure of the BA.2.75 S-trimer in complex with S309 at 3.5 Å, together with structures of the *apo* BA.2.75 and previously reported BA.1-S309 complex, allowing us to dissect the detailed structural variations (Fig. S5A and S5B). Upon S309 binding, BA.2.75 displays a pivotal conformational alteration in the hairpin loop (residues 366-377), separating α1 and α2 helices away, akin to structural arrangement in *apo* BA.1 and BA.1-S309 complex (Fig. 5D and S5C). The mutation D339H lost one hydrogen bond that would be established by Y100 from HCDR3 in BA.1-S309 complex, but established more hydrophobic interactions with Y100 and L110 from S309 in BA.2.75, slightly increasing its binding affinity to BA.2.75 compared to BA.1 (Fig. 5E). These suggest Sotrovimab may become active against BA.2.75, which requires authentic virus validations.

### BA.2.75 displays distinct RBD antibody evasion patterns from BA.4/BA.5

To further investigate the antibody escaping mechanism of BA.2.75, we determined the neutralizing activities against BA.2.75, BA.2.76, and BA.4/BA.5 of a panel of BA.2-effective NAbs from five epitope groups, defined by unsupervised clustering based on high-throughput yeast-display-based deep mutational scanning (DMS), that could be potentially affected by D339H, N460K, R493Q and G446S ^4,26,30^ (Table S3). BA.2.75 can cause a global reduction in neutralizing activities of Group A NAbs, represented by DXP-604 and P2C-1F11 (BRII-196) (Fig. 6A) ^24,31,32^. N460K is the major mutation that caused this antibody evasion. Analyses of the representative structure of P2C-1F11 (BRII-196) suggest K460 with a longer side chain could induce steric clashes that result in NAb evasions (Fig. 6A) ^32^. Interestingly, R493Q can cause neutralization reduction of Group A NAbs isolated from individuals infected by BA.1, which carries R493, while the reversion can also induce neutralization recovery of Group A NAbs isolated from individuals stimulated by wildtype (WT) SARS-CoV-2, which carries Q493 (Fig. 6A). Importantly, although BA.4/BA.5 could induce neutralization reduction of Group A NAbs, BA.2.75 causes more severe Group A antibody evasion, especially for those NAbs that remain highly effective against BA.4/BA.5 (Table S3).

**Fig. 6.**
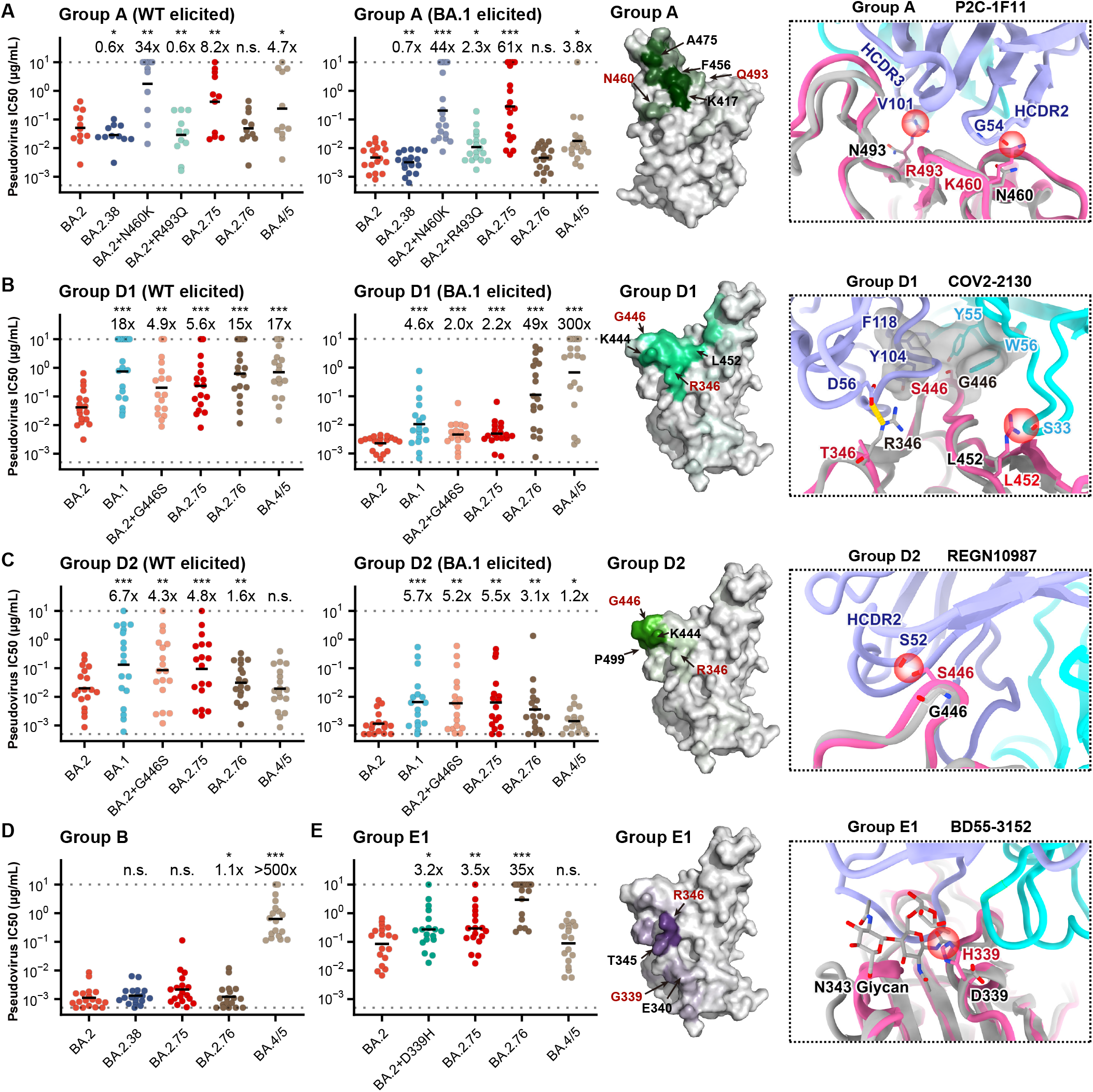
Evasion of NAbs targeting various RBD epitopes by BA.2.38, BA.2.75 and BA.2.76. (A-E) Neutralizing activities against SARS-CoV-2 Omicron variants of NAbs in (A) Group A (N=11 isolated from SARS-CoV-2WT convalescents or vaccinees; N=18 from post-vaccination BA.1 convalescents), (B) Group D1 (N=17, 18, respectively), (C) Group D2 (N=18, 17, respectively), (D) Group B (N=18), and (E) Group E1 (N=18). Deep mutational scanning (DMS) profiles of antibodies in Group A, D1, D2, E1 were projected onto RBD structure to show interacting hotspots of each group. Color shades indicate escape scores of RBD residues. Interface structural models of representative antibodies in Group A, D1, D2, and E1, in complex of RBD, show potential escaping mechanism of BA.2.75 and BA.2.76. Geometric mean of IC50 fold changes compared to BA.2 are annotated above the bars. *p < 0.05; **p < 0.01; ***p<0.001. P-values were calculated using two-tailed Wilcoxon singed-rand test of paired samples.

Reduced activities against BA.2.75 were also observed for NAbs in Groups D1 and D2, mostly due to G446S via disruption of the hydrophobic interactions, conferring the neutralizing activities of these NAbs against BA.2.75 are quite similar to those against BA.1 (Fig. 6B and 6C). Due to the L452R mutation, BA.4/BA.5 could heavily reduce the potency of D1 antibodies (e.g. COV2-2130, a representative antibody from Group D1). Similarly, the R346T mutation, carried by BA.2.76, could also decrease the neutralization activity of Group D1 NAbs. Notably, D1 NAbs isolated from BA.1 convalescents exhibit stronger resistance to BA.2.75, and overall, D1 NAbs are more vulnerable to BA.4/BA.5 compared to BA.2.75; however, this property is not observed in Group D2 NAbs, where BA.2.75 confers more neutralization reduction than BA.4/BA.5, since D2 NAbs are not sensitive to the L452 mutations, but to G446S (e.g. REGN10987, a representative antibody from Group D2) ^4,33^.

Furthermore, Group B antibodies were not affected by BA.2.75, but most of them were completely escaped by BA.4/BA.5 due to F486V, as reported previously (Fig. 6D). D339H, a new specific mutation observed in BA.2.75, could cause an overall neutralization reduction for BA.4/5 effective NAbs in Group E1, which could be largely attributed to the altered configuration of N343 glycan (e.g. BD55-3152, a representative antibody from Group E1) (Fig. 6E) ^4^. Together, the results suggest all four additional RBD mutations of BA.2.75 compared to BA.2 are capable of escaping several groups of NAbs, and the RBD antigenicity of BA.2.75 is distinct from that of BA.4/BA.5, which may partially explain BA.2.75’s strong humoral immune evasion observed in the convalescent plasma from BA.5 breakthrough infection.

### BA.2.75 evades BA.5 effective anti-NTD NAbs

Besides mutations on RBD, BA.2.75 also harbors multiple NTD mutations that may cause anti-NTD NAb evasion ^28^. We tested in total 25 NTD-directing NAbs of their neutralizing capability against SARS-CoV-2 VOCs, including BA.2.75 and BA.2.76 (Table S3 and S6A). Interestingly, anti-NTD NAbs displayed diverse reactivity against Omicron variants (Fig. 7A). A proportion of these NAbs, such as XG2v-024, could broadly neutralize all VOCs, including BA.2.75, with good potency; while some NAbs could effectively neutralize BA.2 and BA.5 but showed decreased activity against BA.2.75 and BA.2.76. (Fig. 7A and S6A). This suggests that NTD targeting NAbs also consists of diverse epitopes and reacts differently to the mutations harbored by Omicron subvariants. To dissect this nature, we first examined the cryo-EM structure of XG2v-024 in complex with BA.2.75 spike (Fig. S7). XG2v024/Spike^BA.2.75^ reveals only one configuration: three XG2v024 Fabs bound to a completely closed S-trimer with three RBDs in the down state (Fig. 7B). This is contrary to the two conformations observed in the *apo* BA.2.75 S-trimer, indicative of a role of XG2v024 in allosteric modulation on RBD “down” disposition. Interestingly, the binding of XG2v024 created steric clashes with the adjacent “up” RBD or its bound hACE2, which reveals the neutralization mechanism via distal RBD conformation modulation and further blockade of ACE2 binding (Fig. 7C). Remarkably, all XG2v024 epitope residues are extremely conserved epitopes across nearly all circulating SARS-CoV-2 variants, enabling XG2v-024’s broad SARS-CoV-2 neutralizing ability (Fig. 7D and Fig. 7E).

**Fig. 7.**
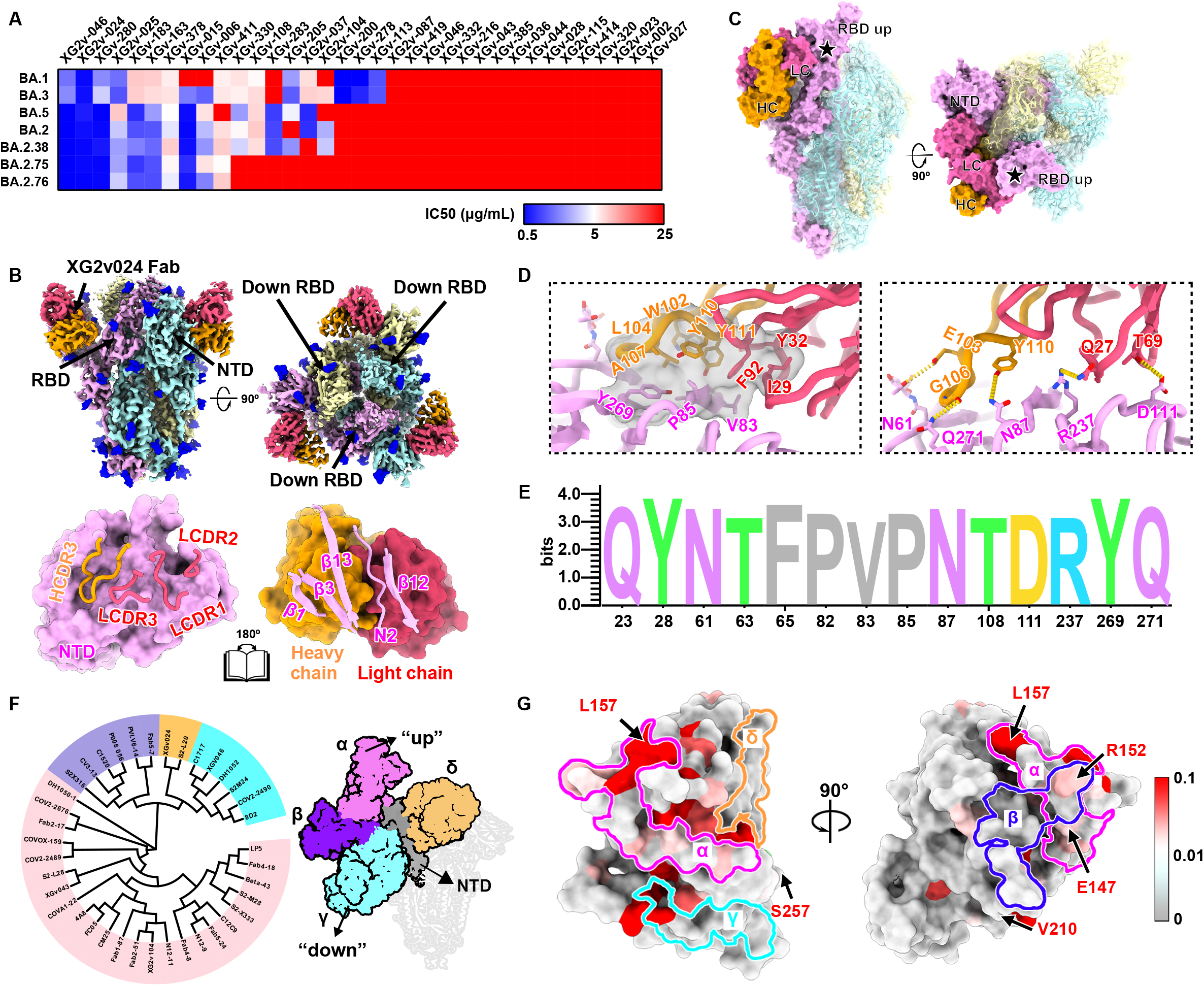
BA.2.75 and BA.2.76 evolve mutations to escape NTD-targeting NAbs. (A) Heatmap of pseudo-typed virus neutralization by antibodies recognized NTD. (B) Cryo-EM structures of the BA.2.75 S-trimer in complex with XG2v024 Fab (top). Structures are shown as surface. The three subunits of S protein are colored in yellow, cyan, and magenta, respectively. The heavy chain and light chain are colored in hot pink and orange, respectively. Interactions between the XG2v024 and BA.2.75 NTD (bottom). The CDRs of the XGv024 that interact with BA.2.75 NTD are displayed as thick tubes over the magenta surface of the NTD. The XG2v024 epitope is shown as a cartoon representation over the surface of the XG2v024 Fab. (C) XG2v024 neutralizing mechanism. The BA.2.75 S-trimer and XG2v024 Fab are shown as surface. The color scheme remains unchanged from the previous panels. Steric clash is marked as black stars. (D). Interactions details of BA.2.75 NTD in complex with XG2v024 Fab. The hydrophobic (top) and hydrophilic (bottom) interactions are shown, respectively. Hydrogen bonds and hydrophobic patches are presented as yellow dashed lines and gray surfaces, respectively. (E)Analysis of sequence conservation of XG2v024 epitope. The logo plot represents the conservation of XG2v024 epitope residues from 26 SARS-CoV-2 lineages. (F)Structure-based antigenic clustering of SARS-CoV-2 NTD Nabs (left). A total of 38 NTD NAbs with available structures were classified into four clusters (α, β, γ and δ). Surface representative model of four types of NAbs bound to the NTD (right). Fab fragments of four representative antibody are shown in different colors and the NTD is colored in gray. (G) Structural landscapes of the four classes of NTD NAbs. Antigenic patches recognized by four types of NAbs are outlined in the assigned color scheme. Five mutations in the BA.2.75 NTD are labeled.

By further examining the available structures of 38 SARS-CoV-2 NTD-targeting NAbs, including XG2v-024 presented here, we found that the NTD NAbs can be classified into four classes and the NTD mutations of BA.2.75 affects different classes (Fig. 7F and 7G). Class α antibodies, targeting the NTD super-site and facing away from the viral membrane (facing up), possess limited neutralizing breadth due to highly frequent mutations on the epitopes (Fig. S6B and S6C) ^15,34^. Sites β and δ (e.g., targeted by XGv024), as the left and right flank clusters, construct a shallow groove and locate at the back of the groove, respectively, eliciting relatively broad neutralizing antibodies, albeit with less potency (Fig. 7F). Importantly, the new NTD mutations K147E, W152R, F157L, and G257S carried by BA.2.75 and the Y248N carried by BA.2.76 are located on the epitope of Class α and β antibodies, likely causing the escape of those BA.5 effective NTD NAbs. By contrast, γ antibodies bound to a patch beneath the groove have their Fab constant domains directed downward toward the virus membrane (facing down) and were proved to enhance infection efficiency *in vitro*; unfortunately, the V213G harbored by all Omicron variants could cause large-scale escapes of these NAbs (Fig. 7F and S6C). Noteworthy, the classification is highly similar to the classes assigned in biolayer interferometry competition experiments ^34^. Together, we showed that besides the RBD mutations, the new NTD mutations evolved by BA.2.75 could also induce further neutralization escape of BA.2 and BA.5 effective anti-NTD NAbs.

## Discussion

The rapid evolution of SARS-CoV-2 Omicron variants greatly challenged the efficacy of vaccines and therapeutics ^3,35–39^. In this paper, we demonstrated that the local growth advantage of BA.2.75 compared to BA.5 could be explained by the previous pandemic caused by Delta VOC in India ^40^. As Delta and BA.4/5 both harbor L452R mutation on the RBD, and arginine generally being an immunogenic residue, convalescent plasma from Delta infection may contain 452R-targeting BA.5-effective NAbs, which could impair the transmissibility of BA.4/BA.5.

More importantly, we showed that the BA.5 convalescent plasma could not neutralize BA.2.75 well, suggesting that BA.2.75 could potentially become the next dominant strain after the global BA.4/5 wave. Also, it is worrisome that the significantly enhanced hACE2-binding affinity of BA.2.75 could lead to its higher capability to tolerate additional immune-evasive mutations, which usually results in the reduction of ACE2-binding. Indeed, several mutations that were proved antibody-evasive, such as R346T and L452R, have been identified in sublineages of BA.2.75, which would further increase its humoral evasion capability and transmission advantage. Our results urge the close monitoring of the spread of BA.2.75.

## Acknowledgments

This project is financially supported by the Ministry of Science and Technology of China and Changping Laboratory under the project number (CPL-1233).

## Author contributions

Y.C., X.S.X and X.W. designed the study. Y.C. and X.W. and wrote the manuscript with inputs from all authors. F.J. and J.W. (BIOPIC) performed and analyzed the yeast-display-based deep mutational scanning experiments. Y.C., W.S., A.Y., Y.Y., T.X., P.W., J.W. (Changping Laboratory), S.Y., L.L., R.A., Yao W., N.Z., X.N., L.Y., C.Y., F.S. performed and analyzed the pseudovirus neutralization assays. Y.Y. and Youchun W. prepared the VSV-based SARS-CoV-2 pseudovirus. L.W., P.L., C.Y., Q.Z., J.D., W.F., and X.W. performed the structural analyses. Q.G. proofed the manuscript. C.Y., R.F. and X.W. performed surface plasma resonance (SPR) experiments. X.H. and R.J. recruited the SARS-CoV-2 vaccinees and Omicron/Delta convalescents.

## Methods

### Protein expression and purification

The full-length Spike (S) proteins of Omicron (BA.1) and its variants BA.2, BA2.12.1, BA.4/5 were derived from previous constructions. BA.2 RBD and its mutation N460K, BA.2.75 spike (T19I, L24S, Δ25-27, G142D, K147E, W152R, F157L, I210V, V213G, G257S, G339H, S371F, S373P, S375F, T376A, D405N, R408S, K417N, N440K, G446S, N460K, S477N, T478K, E484A, Q498R, N501Y, Y505H, D614G, H655Y, N679K, P681H, N764K, D796Y, Q954H, N969K) and BA.2.75 RBD and its mutations H339D, S446G, K460N, Q493R were realized by overlapping PCR with BA.2 Spike gene as template. To facilitate protein expression and stabilize the trimer conformation, all S gene constructs have proline substitutions at residues 817, 892, 899, 942, 986, 987 and alanine substitutions at residues 683, 685, and the T4 fibrin fold domain added to the C-terminal target sequence (ACRO Biosystems, cat. no. SPN-C5523). In addition, His or Strep II tags were connected to the C-terminal of spike and RBD sequences to facilitate protein purification. For the selected antibody, the light and heavy chains need to be constructed separately. All the constructed plasmids were transiently transfected into suspended HEK293 F cells and cultured in a constant temperature shaker of 8% CO_2_ and 37 °C. After 72 hours of culture(Antibodies require 120 hours), cell supernatants were collected and Spike and RBD were initially purified by affinity chromatography using Ni-NTA or StrepTactin resin, while antibodies required protein A purification. The proteins were repurified using Superdex 200 10/300GL (Cytiva) or Superose 6 10/300 (Cytiva) in 20 mM Tris-HCl, 200 mM NaCl, pH 8.0.

### Determination of thermal stability

The thermofluor stability assay was performed to evaluate the stability of Spike and RBD of omicron and its variants BA.1, BA.2, BA.2.12.1, BA.2.75 and BA.5(ACRO Biosystems). For the Spike protein, we performed two different pH conditions (pH = 7.4 and pH = 5.5), while for the RBD, only stability assay was performed at pH = 7.4. All protein samples were set up as 25 uL reaction system, including 5 mg protein and 5000× SYPRO Orange (Invitrogen, Carlsbad, USA) as fluorescence probe. The MX3005 qPCR instrument (Agilent, Santa Clara, USA) was used to detect the fluorescent signal generated by the protein during heating from 25°C to 99°C at a rate of 1°C/min. GraphPad Prism 9.0.1 (GraphPad Software Inc.) was used to draw the temperature curve.

### Bio-layer interferometry

Bio-layer interferometry (BLI) experiments were performed on an Octet Red 96 instrument (Fortebio) to measure the binding affinities of S309 NAb. S309 was immobilized onto Protein A biosensors (Fortebio). BA.1 RBD, BA.2 RBD, BA.2.12.1 RBD, BA.2.75 RBD and BA.4/5 RBD and in PBS used as analytes were diluted by threefold serial dilutions. The Octet BLI Analysis 9.1 (Fortebio) software was used to analyze the experimental data using a 1:1 fitted model.

### Fab generation

Fab fragments were prepared using the Pierce Fab Preparation Kit (Thermo Scientific) as previously described ^41^. Briefly, the sample first needs to be subjected to a desalting column to remove salts. Then, the effluent was collected and incubated with papain attached to the beads. The Fab fragments were cleaved from the antibody for 5 hours at 37 °C. The mixture was then transferred to a Protein A column to purify the Fab fragments (ThermoFisher, Catalog (Cat.) No.). 10010023).

### MD stimulation and RMSF calculating

Initial models of BA.2 and BA.2.75 RBD were from the structure 7XIW downloading from PDB and the structure of apo S-Trimer of BA.2.75 at neutral pH determined by this study, respectively. Before final stimulation, we used CHARMM-GUI to generate the inputs for simulation packages GROMACS. After PDB checking, waterbox size specifying, water model specifying (TIP3P), ions adding, periodic boundary condition setting and force fielding specifying (OPLS-AA/M), the data generated was submitted to GROMACS-2021 to Energy Minimization, NVT Equilibration, NPT Equilibration and 10 ns MD simulation. NVT ensemb1e via the Nose-Hoover method at 300 K and NPT ensemble at 1 bar with the Parinello-Rahman algorithm were employed to make the temperature and the pressure equilibrated, respectively. The last 2 ns frames were extracted to calculated RMSF.

### Surface plasmon resonance

Surface plasmon resonance (SPR) experiments were performed on the Biacore 8K (GE Healthcare). Immobilization of human ACE2 on a CM5 sensor chip and injection of purified Omicron and its variant RBDs with its corresponding single point mutation. The response units were recorded by Biacore 8K Evaluation Software (GE Healthcare) at room temperature, and the raw data curves were fitted to a 1:1 binding model using Biacore 8K Evaluation Software (GE Healthcare).

### Cryo-EM sample preparation and data collection

3 μL of purified SARS-CoV-2 BA.2.75 variant Spike trimer (pH 7.4 and pH5.5) at 1.2 mg/mL and BA.2.75 S protein mixed with S309 Fab, XG2v024 Fab and human ACE2 at 1.0 mg/mL in purification buffer solution (20 mM Tris, 200 mM NaCl, pH 7.4) were dropped onto the pre-glow-discharged holey carbon-coated gold grid (C-flat, 300-mesh, 1.2/1.3, Protochips In.) Grids were blotted for 6 s in 100% relative humidity and room temperature for plunge-freezing (Vitrobot; FEI) in liquid ethane. Cryo-EM data sets were collected at a 300 kV Titan Krios microscope (Thermo Fisher) equipped with a K2 or K3 detector. Movies (32 frames, total dose 60 e −Å −2) were recorded with a defocus of from −1.5 to −2.7 μm using SerialEM, which yields a final pixel size of 1.07 Å or 1.04 Å.

### Data processing and model refinement

A total of 6,459 micrographs of pH 7.4 BA.2.75 S protein, 4,886 micrographs of pH 5.5 BA.2.75 S protein, 5,761 micrographs of BA.2.75 S in complex with S309 Fab, 5,606 micrographs of BA.2.75 S in complex with XG2v024 and 7,225 micrographs of BA.2.75 S in complex with ACE2 were recorded. The CTF values were estimated by Patch CTF in cryoSPARC (v3.3.2). Particles were picked based on templates and extracted for 2D Classification. High-quality particles were selected and passed to Heterogeneous Refinement for 3D classification. Then, Homogeneous Refinement and Non-uniform Refinement were performed for high-resolution reconstruction. To obtain reliable interface density, Local Refinement was used to further improve the resolution of interfaces between BA.2.75 S and S309 Fab, XG2v024 Fab and ACE2. The atom models were generated by fitting the apo BA.2 S trimer (PDBID:7XIW, 7XIX), S309 Fab (PDBID:7TLY), ACE2 (6M0J), XG2v024 (PDBID:7CHS for heavy chain and 7N4L for light chain) into the cryo-EM density by Chimera. Then the models were adjusted manually in Coot and real-space refinement in Phenix.

### Plasma isolation

Blood samples were obtained from SARS-CoV-2 vaccinee convalescent individuals who had been infected with Delta, BA.1, BA.2, and BA.5. Written informed consent was obtained from each participant in accordance with the Declaration of Helsinki. Whole blood samples were diluted 1:1 with PBS+2% FBS and then subjected to Ficoll (Cytiva, 17-1440-03) gradient centrifugation. After centrifugation, plasma was collected from the upper layer. Plasma samples were aliquoted and stored at −20 °C or less and were heat-inactivated before experiments.

### Pseudovirus neutralization assay

SARS-CoV-2 variants Spike pseudovirus was prepared based on a vesicular stomatitis virus (VSV) pseudovirus packaging system. D614G spike (SARS-CoV-2 spike (GenBank: MN908947) +D614G), BA.1 spike (A67V, H69del, V70del, T95I, 142-144del, Y145D, N211del, L212I, ins214EPE, G339D, S371L, S373P, S375F, K417N, N440K, G446S, S477N, T478K, E484A, Q493R, G496S, Q498R, N501Y, Y505H, T547K, D614G, H655Y, N679K, P681H, N764K, D796Y, N856K, Q954H, N969K, L981F), BA.2 spike (GISAID: EPI_ISL_7580387, T19I, del24-26, A27S, G142D, V213G, G339D, S371F, S373P, S375F, T376A, D405N, R408S, K417N, N440K, S477N, T478K, E484A, Q493R, Q498R, N501Y, Y505H, D614G, H655Y, N679K, P681H, N764K, D796Y, Q954H, N969K), BA.2.38 spike (BA.2+N417T), BA.2.12.1 spike (BA.2+L452Q+S704L), BA.2.75 spike (BA.2+K147E+W152R+F157L+I210V+G257S+D339H+G446S+N460K+R493Q), BA.2.76 spike (BA.2+R346T+Y248N), BA.4/BA.5 spike (T19I, L24S, del25-27, del69-70, G142D, V213G, G339D, S371F, S373P, S375F, T376A, D405N, R408S, K417N, N440K, L452R, S477N, T478K, E484A, F486V, Q498R, N501Y, Y505H, D614G, H655Y, N679K, P681H, N764K, D796Y, Q954H, N969K) plasmid is constructed into pcDNA3.1 vector. G*ΔG-VSV virus (VSV G pseudotyped virus, Kerafast) and spike protein plasmid were transfected to 293T cells (American Type Culture Collection [ATCC], CRL-3216). After culture, the pseudovirus in the supernatant was harvested, filtered, aliquoted, and frozen at −80°C for further use.

Huh-7 cell line (Japanese Collection of Research Bioresources [JCRB], 0403) was used in pseudovirus neutralization assays. Plasma samples or antibodies were serially diluted in culture media and mixed with pseudovirus, and incubated for 1 h in a 37°C incubator with 5% CO_2_. Digested Huh-7 cells were seeded in the antibody-virus mixture. After one day of culture in the incubator, the supernatant was discarded. D-luciferin reagent (PerkinElmer, 6066769) was added into the plates and incubated in darkness for 2 min, and cell lysis was transferred to the detection plates. The luminescence value was detected with a microplate spectrophotometer (PerkinElmer, HH3400). IC50 was determined by a four-parameter logistic regression model.

---

**Table S1.**Reports of detailed parameters of the reconstructed Cryo-EM structures.

**Table S2.** Summarized information of SARS-CoV-2 vaccinated individuals and convalescents involved in the study.

**Table S3.** Summarized information and experiment results of SARS-CoV-2 neutralizing antibodies involved in the study.

**Figure S1.**
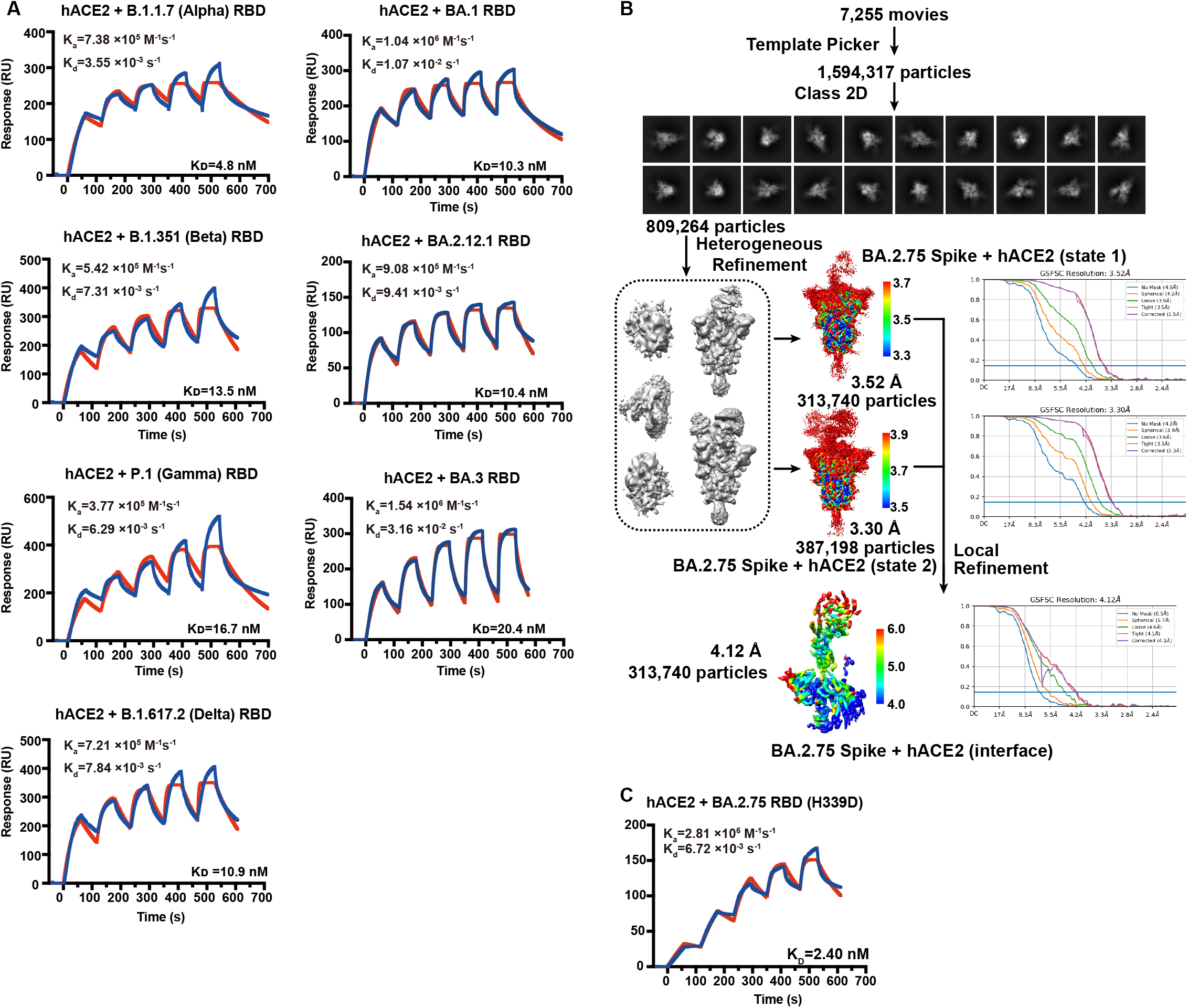

**Figure S2.**
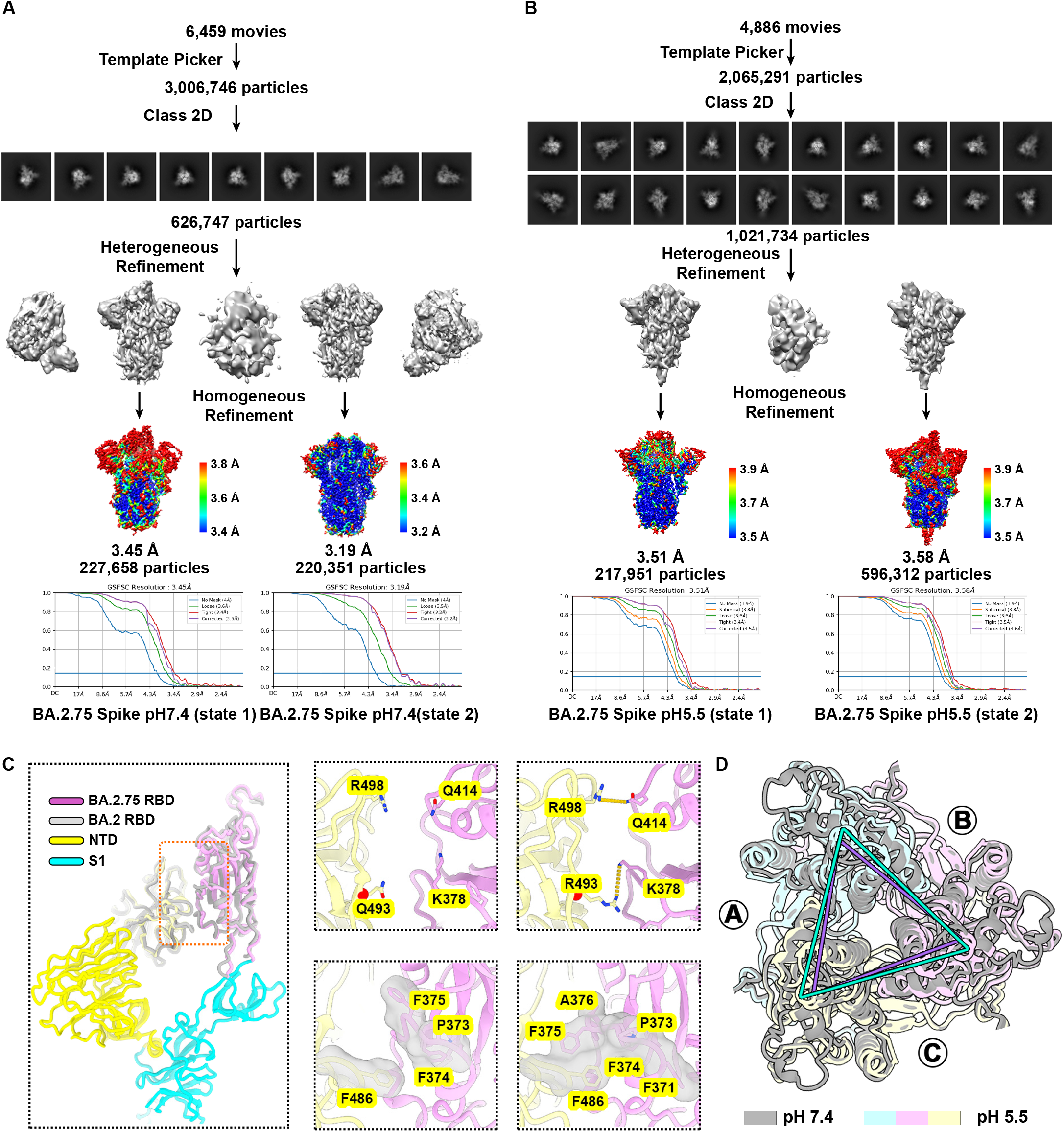
Structural analyses of BA.2.75 spike in different pH, Related to Figure 3. (A-B) Flow charts for cryo-EM data processing of BA.2.75 spike at (A) neutral pH and (B) acidic pH. (C) Structural features underpinning the up configuration. Left: Cartoon representation of BA.2.75 and BA.2 S-trimer in a prefusion conformation with one protomer in “open” state. The BA.2.75 RBD and BA.2 RBD are colored in magenta and gray, respectively. The NTD domain and S2 domain are colored in yellow and cyan, respectively. The zoomed-in view of interaction details of two independent interfaces for BA.2.75 (middle panel) and BA.2 (right panel). The mutated residues are shown as sphere in red, and the residues involved in the interactions are shown as sticks. The hydrogen bonds are shown as yellow dashed lines and hydrophobic network is highlighted in gray. (D). Superimposition of S2 subunit of the BA.2.75 S-trimer (pH7.4, gray) onto the BA.2.75 S-trimer (pH5.5, yellow, lightblue, pink).

**Figure S3.**
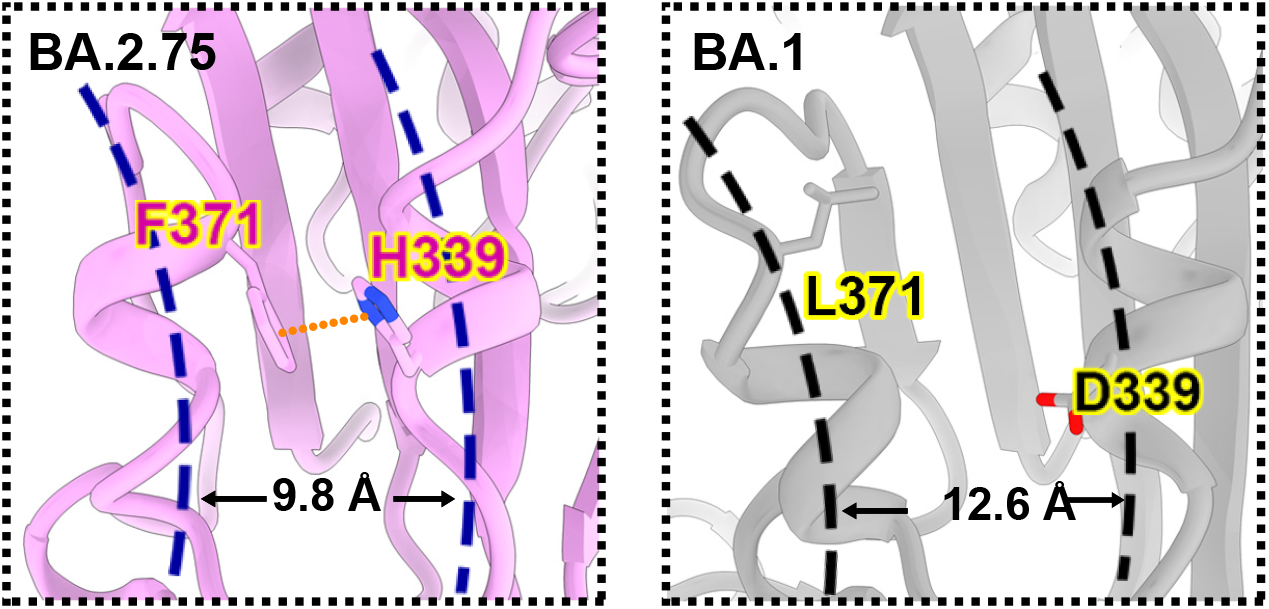
Local conformational change of BA.2.75 RBD compared to BA.1, Related to Figure 4. Structural comparison of α1 and α2 helices on RBDs of BA.2.75 (left) and BA.1 (right). The π-π stack formed between H339 and F371 in BA.2.75 RBD are marked as yellow dashed lines. The distances between α1 and α2 helices on RBD are also highlighted.

**Figure S4.**
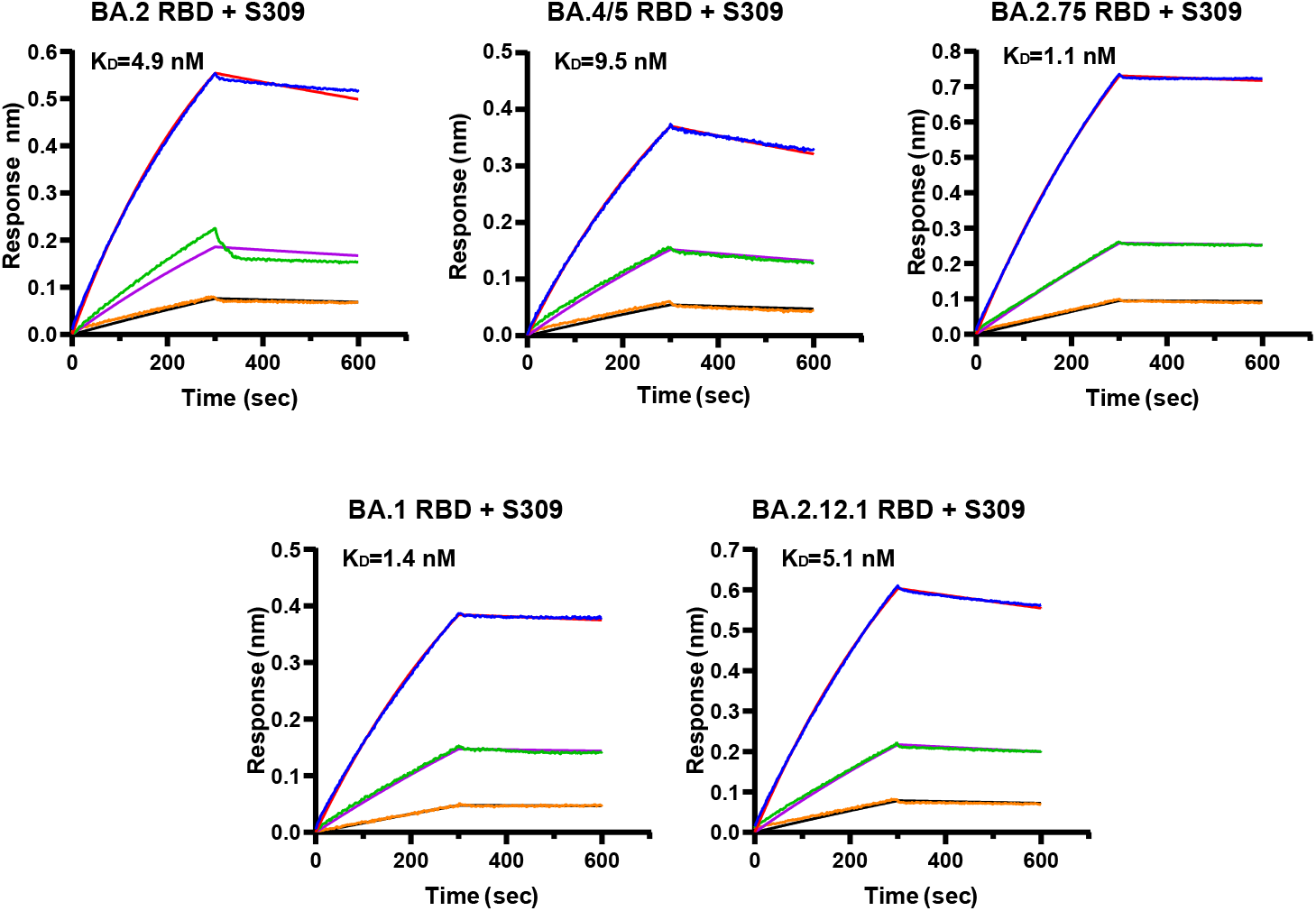
Binding affinity of S309 against Omicron variants RBD, Related to Figure 5. BLI sensorgrams measuring the binding affinity of S309 with BA.1 RBD, BA.2 RBD, BA.2.12.1 RBD, BA.4/5 RBD and BA.2.75 RBD.

**Figure S5.**
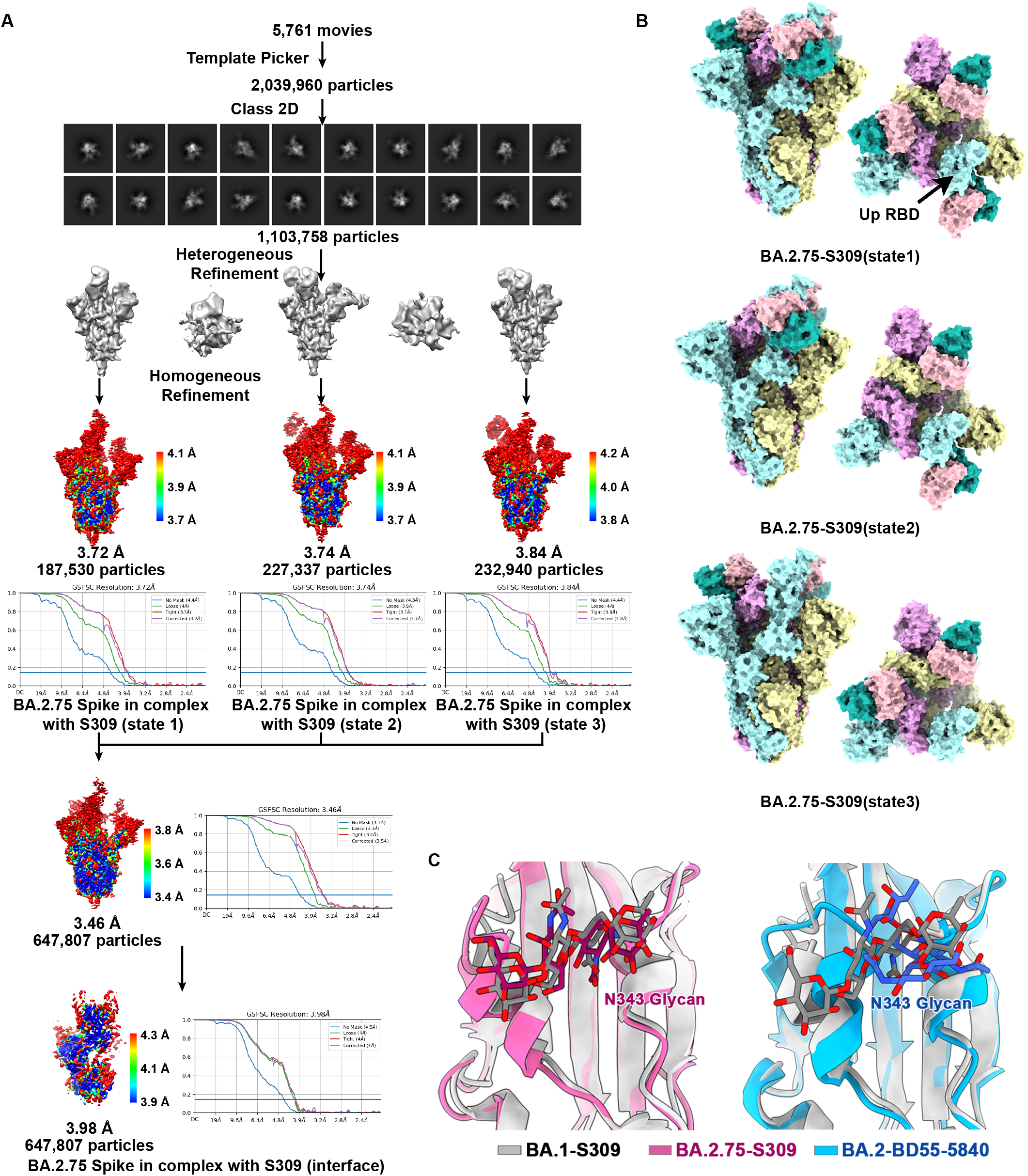
Structural analyses of BA.2.75-S309 complex, Related to Figure 5. (A) Flow charts for cryo-EM data processing of BA.2.75 spike and S309 complex. (B) Surface presentations of the three states of BA.2.75 S-trimer in complex with S309 Fab. The three subunits of S protein are colored in yellow, cyan, and magenta, respectively. The heavy chain and light chain are colored in light pink and light seagreen, respectively. (C) Diagram presentation of N343 glycan conformational differences among BA.1 bound to S309 (gray), BA.2 bound to S309 (pink), and BA.2 bound to BD55-5840 (blue) are shown.

**Figure S6.**
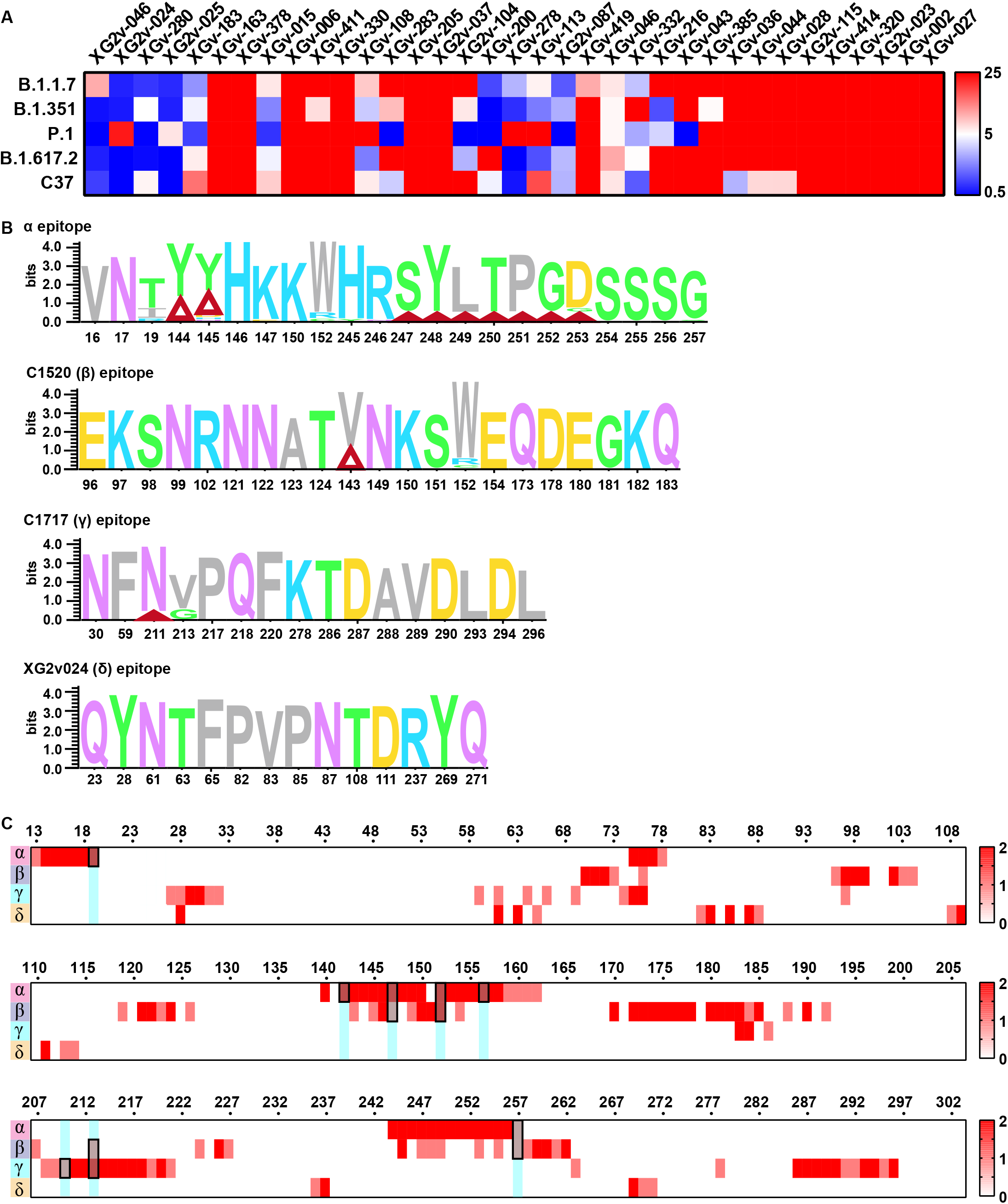
Characteristics of NTD-targeting antibodies, Related to Figure 7. (A) Heatmap of pseudo-typed virus neutralization by antibodies recognized NTD (unit: μg/ml). (B) Analysis of sequence conservation of four classes (α, β, γ and δ) epitopes of NTD recognized antibodies. The logo plot represents the conservation of epitopes residues from 26 SARS-CoV-2 lineages: WT, Alpha, Beta, Gamma, Lambda, Mu, Delta, Delta plus, BA.1, BA.1.1, BA.2, BA.2.12.1, BA.2.13, BA.4, BA.5, BA.2.75, Eta, Lota, Kappa, Theta, Iota, B.1.1.318, B.1.620, C.1.2, C.363 and Epsilon. Deletions on NTD are represented as red triangles. (C) Heatmap represents the frequency of NTD residues recognized by NAbs from four classes (α, β, γ and δ). Mutations present in BA.2.75 NTD are marked out and highlighted.

**Figure S7.**
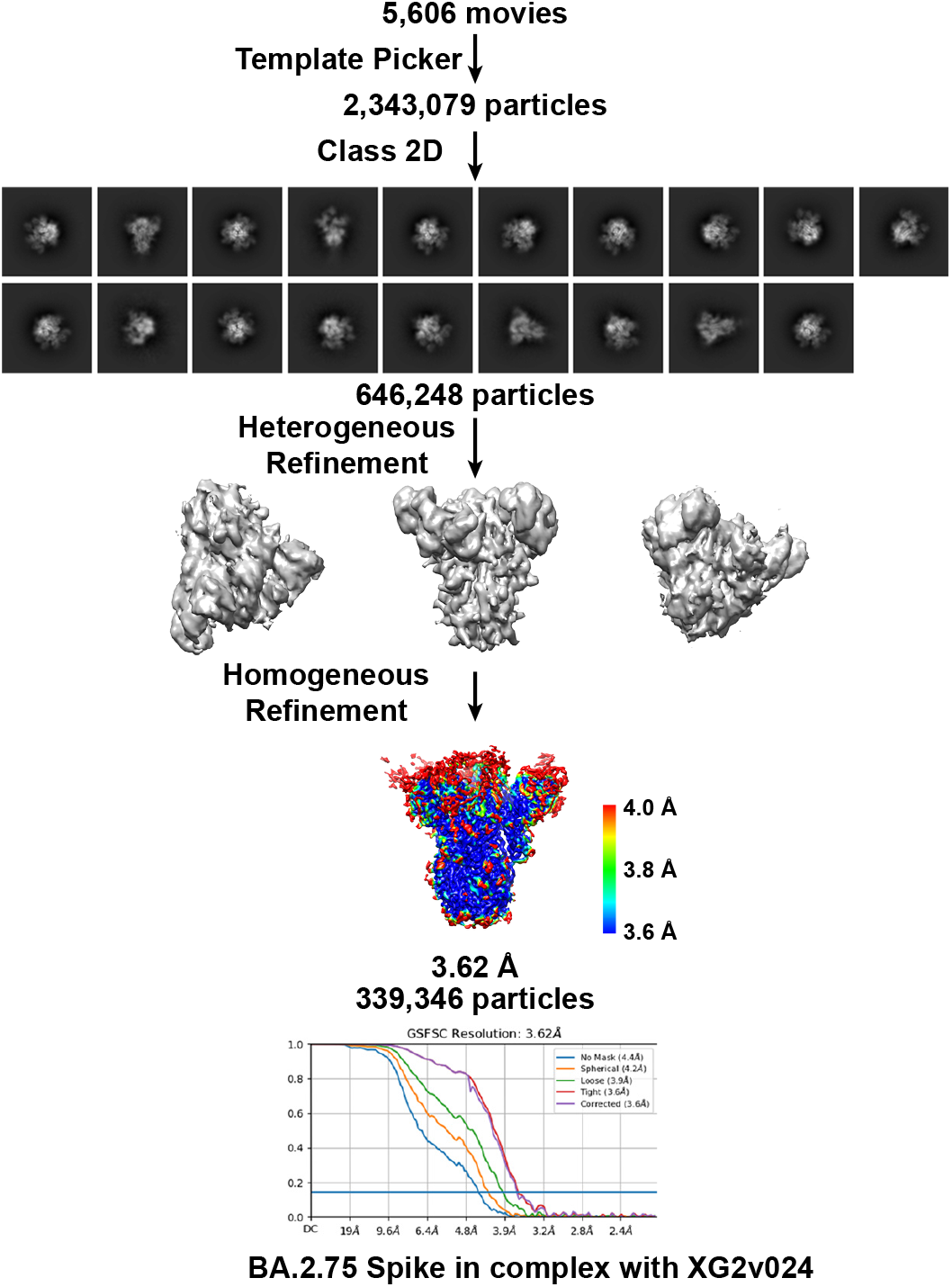
Cryo-EM structures of BA.2.75 spike in complex of XG2v024, Related to Figure 7. Flow charts for cryo-EM data processing of BA.2.75 spike and XG2v024 complex.

